# More similarity than difference: comparison of within- and between-sex variance in early adolescent brain structure

**DOI:** 10.1101/2024.08.15.608129

**Authors:** Carinna Torgerson, Kathserine Bottenhorn, Hedyeh Ahmadi, Jeiran Choupan, Megan M. Herting

## Abstract

**Background:** Adolescent neuroimaging studies of sex differences in the human brain predominantly examine mean differences between males and females. This focus on between-groups differences without probing relative distributions and similarities may contribute to both conflation and overestimation of sex differences and sexual dimorphism in the developing human brain.

**Methods:** We aimed to characterize the variance in brain macro-and micro-structure in early adolescence as it pertains to sex at birth using a large sample of 9-11 year-olds from the Adolescent Brain Cognitive Development (ABCD) Study (N=7,723). Specifically, for global and regional estimates of gray and white matter volume, cortical thickness, and white matter microstructure (i.e., fractional anisotropy and mean diffusivity), we examined: within-and between-sex variance, overlap between male and female distributions, inhomogeneity of variance via the Fligner-Killeen test, and an analysis of similarities (ANOSIM). For completeness, we examined these sex differences using both uncorrected (raw) brain estimates and residualized brain estimates after using mixed-effects modeling to account for age, pubertal development, socioeconomic status, race, ethnicity, MRI scanner manufacturer, and total brain volume, where applicable.

**Results:** The overlap between male and female distributions was universally greater than the difference (overlap coefficient range: 0.585 -0.985) and the ratio of within-sex and between-sex differences was similar (ANOSIM R range: -0.001 -0.117). All cortical and subcortical volumes showed significant inhomogeneity of variance, whereas a minority of brain regions showed significant sex differences in variance for cortical thickness, white matter volume, fractional anisotropy, and mean diffusivity. Inhomogeneity of variance was reduced after accounting for other sources of variance. Overlap coefficients were larger and ANOSIM R values were smaller for residualized outcomes, indicating greater within-and smaller between-sex differences once accounting for other covariates.

**Conclusions:** Reported sex differences in early adolescent human brain structure may be driven by disparities in variance, rather than binary, sex-based phenotypes. Contrary to the popular view of the brain as sexually dimorphic, we found more similarity than difference between sexes in all global and regional measurements of brain structure examined. This study builds upon previous findings illustrating the importance of considering variance when examining sex differences in brain structure.

**Highlights:**

- High male/female overlap is ubiquitous across all brain features in early adolescence
- Male variance exceeded female variance for global and regional brain volumes
- Between-and within-sex differences were similar in magnitude for all features

**Plain English Summary:** Brain imaging research has consistently revealed differences between males and females in the shape and size of adolescent brains. Studies usually compare the average male brain to the average female brain. However, brain structure varies greatly among individuals, even within the same sex. Without looking at both the variability within people of the same sex, and the degree of similarity between the sexes, it is unclear if separating adolescent brains into male and female categories will help us understand brain development. In this study, we looked at the overlap in brain structure among male and female youths (ages 9 to 11 years). We also compared variability between sexes and within each sex. Overall, we found that, there was more similarity than difference between male and female brains. The difference between any given male and any given female was similar to the difference between two individuals of the same sex. These findings suggest that, despite some small average differences, the brains of early adolescent males and females are more alike than different at ages 9-11 years.

## Background

Sexual dimorphism refers to traits with two distinct forms, each existing predominantly or exclusively among one sex, whereas sex differences describe traits that fall along a continuum, but exhibit a difference in mean or variability between males and females (DeCasien et al., 2022; McCarthy et al., 2012). In the neuroscience literature, the conflation of the terms is exacerbated by researchers’ tendency to focus on mean sex differences. For example, when interpreting sex differences, the mean trait or phenotype is often generalized to the entire sex (i.e., “males have larger brains than females”) (Sanchis-Segura et al., 2022). In addition to differences attributable to differential expression of X-and Y-chromosome genes, the organizational-activational hypothesis posits that sex differences in exposure to steroid hormones during puberty cause both structural and functional sex differences in the brain and other non-gonadal tissues (A. P. Arnold, 2009; McCarthy et al., 2009; K. M. Schulz et al., 2009). This makes adolescence a crucial period of study for the development of sex differences in the brain.

Adolescent studies of sex differences in brain structure predominantly test for significant mean group differences between males and females (Giedd et al., 2012; Giedd & Denker, 2015; Kaczkurkin et al., 2019; Lenroot & Giedd, 2010). On average, regional cortical volumes are larger among male adolescents than among female adolescents (Gennatas et al., 2017; Paus et al., 2010), as are a number of subcortical regions, including the putamen, pallidum, amygdala, thalamus, and cerebellum (Adeli et al., 2020; Paus, 2010; Paus et al., 2010). However, some authors have reported greater whole-brain cortical thickness in adolescent females than in males (Zhou et al., 2015), while others reported no sex differences (Bramen et al., 2012; Menary et al., 2013; Vijayakumar et al., 2016). In addition to increased gray matter volume, male adolescents also display increased white matter volumes relative to female adolescents (Pfefferbaum et al., 2016). Studies of fractional anisotropy (FA) and mean diffusivity (MD) -measures of white matter microstructure commonly used to study white matter development and integrity – have shown mixed results. For example, some studies report higher FA in male adolescents compared to females (Herting et al., 2012a; Lawrence et al., 2023; Pohl et al., 2016; Torgerson et al., 2024) while others report higher FA in female adolescents (Bava et al., 2011; Schmithorst et al., 2007). However, females enter puberty and reach maturity at younger ages than males (Brix et al., 2019). Similarly, measures of gray and white matter structure peak earlier in girls than in boys (Raznahan et al., 2011a; Simmonds et al., 2014). Therefore, it is important to account for differences in both maturation and chronological age when studying peripubertal development.

Despite relatively small effect sizes, numerous studies have concluded that these differences amount to sexual dimorphism in the developing brain (Brennan et al., 2021; Herting et al., 2012b; Lenroot et al., 2007; Paus et al., 2010; Seunarine et al., 2016; Yang et al., 2021). This elevation of sex differences to sexual dimorphism inappropriately uses aggregate statistical results to infer the nature of inter-individual relationships, which is a form of ecological fallacy (Gnaldi et al., 2018; Nieri et al., 2003; Paik, 1985). For example, though males have - on average – 9–10% larger brains in adolescence (Giedd et al., 1997, 2015; Lenroot et al., 2007), this statistic alone does not indicate that a randomly selected female is more likely than not to have a regional brain volume below a randomly selected male or below the population mean. Similarly, a mean sex difference is not sufficient evidence to claim that all females are more similar to each other than to any males. Such a comparison would require a deeper understanding of the dispersion of the data, particularly the relative within-and between-sex variance (Warton & Hui, 2017). Therefore, more nuanced statistical approaches are required to more fully contextualize the sex differences noted in the existing neuroimaging literature. In fact, in adults, overlap distribution statistics and formal analyses of similarity have shown extensive overlap between the distributions of MRI brain outcomes for each sex (N = 1,403; total age ranges 12-75 years) (Joel et al., 2015) and that brain metrics from two random individuals of the same sex differ as much as those from two random individuals of the opposite sex (Sanchis-Segura et al., 2022). These innovative statistical approaches challenge the narrative of “hard-wired” differences between “male brains” and “female brains” (Amen, 2013; Baron-Cohen, 2009; Blum, 1998; Brizendine, 2006, 2022; Darlington, 2009; Gurian, 2010; Gurian & Stevens, 2006; James, 2009; Lundin, 2009; McKay, 2018; M. L. Schulz, 2005). However, similar research contextualizing sex differences in child and adolescent brains remains sparse.

Using the largest study of brain development - the Adolescent Brain Cognitive Development Study (ABCD Study®) - we recently examined how sex and gender relate to gray matter macrostructure and white matter microstructure in a nationwide U.S. sample of 9-11 year-olds (Torgerson et al., 2024). We found that sex - but not felt- gender - was a significant predictor of early adolescent subcortical volume, cortical thickness, local gyrification index and white matter microstructure in the majority of regions examined. Furthermore, Wierenga, et al. (Wierenga et al., 2018, 2022) previously found that male variability in the volumes of the hippocampus, pallidum, putamen, and cerebral gray and white matter was greater than female variability not only at the sample mean, but also at the extremes upper and lower ends of the distribution for children and adolescents. Building on this work, we examined inter-individual variability in brain development in the ABCD Study and found sex differences in the variability of the annualized percent change (Bottenhorn et al., 2023). Specifically, we reported greater male variability in white matter volumes and network connectivity, but greater female variability in the development of cortical macro-and micro-structure.

Consequently, this study aims to contextualize the cross-sectional relationship between mean group sex differences and inter-individual differences in brain structure in early adolescents ages 9 to 11 years old. Building upon our previous findings showing widespread, yet very small effect sizes (Torgerson et al., 2024), we aimed to further characterize within-and between-group differences, inhomogeneity between the sexes, overlap between the sexes, and conduct an analysis of similarity (ANOSIM) across various macro-and micro-structural brain metrics between males and females. Given large differences in overall head sizes and other potential confounders, we conducted our analyses on both raw (uncorrected) brain metrics and after adjusting for total brain volume (TBV) and other sociodemographic factors. Based on previous studies, we hypothesized that the variance between the male and female means would not exceed the within-sex variance, and that this would be true of more regions after adjusting for covariates. We expected inhomogeneity of variance between males and females, in line with previous research (Bottenhorn et al., 2023; Wierenga et al., 2018). In terms of overlap, we hypothesized that we would find substantial overlap (i.e. greater than 50% overlap) of male and female distributions in all regions and measures examined, and that this overlap would be larger after adjusting for potential confounders, including TBV for volumetric outcomes.

## Methods

### Participants

This study utilized data collected as part of the larger ongoing Adolescent Brain Cognitive Development (ABCD) Study®, which involves 11,880 children at 21 different sites around the United States (*ABCD Study*, 2022; Casey et al., 2018; Hagler et al., 2019). The study included children from diverse geographic, demographic, and socioeconomic backgrounds (Garavan et al., 2018; Heeringa & Berglund, 2020). Children with severe sensory, neurological, medical or intellectual limitations, lack of English proficiency or inability to complete an MRI scan were excluded from the ABCD Study (Li et al., 2021). With respect to age, sex, and household size, the ABCD cohort closely matches the distribution of 9-11-year-olds in the American Community Survey, a large probability sample survey of U.S. households conducted annually by the U.S. Bureau of Census (Heeringa & Berglund, 2020). Raw and minimally processed data are publicly available from the ABCD Study in service of increasing reproducibility. We utilized a combination of raw and tabulated questionnaire and neuroimaging data from the study baseline as obtained from the NDA 3.0 (raw T1 and T2 structural MRI files) and 4.0 (tabulated questionnaire and diffusion MRI) releases (NDA 3.0 and 4.0 data release 2021; https://dx.doi.org/10.15154/1523041). We chose to perform our own preprocessing for gray matter macrostructure using both T1w and T2w images to improve parcellation accuracy (Torgerson et al., 2024).

After obtaining the data, we implemented a series of quality control standards (Supplemental Figure 1). Participants were excluded if their data were collected outside the 21 primary research sites, failed execution of the pre-processing or processing pipelines, failed to meet the raw or post-processing quality control standards of the ABCD consortium (Hagler et al., 2019), or had incidental neurological findings noted by a radiologist (Li et al., 2021). To reduce within-family correlation and meet statistical assumptions for independence, we decided to restrict our sample to one child per family (chosen randomly).

### Sex

The ABCD Study collects parent-reported sex assigned at birth. However, due to the multidimensional nature of sex, assignment at birth is not always an accurate reflection of chromosomal sex. Therefore, we also chose to use the frequency ratio of X and Y alleles to detect the presence of a Y chromosome and ascertain the genetic sex of participants. Children whose assigned sex and genetic sex did not match (n = 9) were excluded from the analysis.

### Neuroimaging Data

A harmonized data collection protocol was utilized across sites with either a Siemens, Phillips, or GE 3T MRI scanner. Motion compliance training, as well as real-time, prospective motion correction was used to reduce motion distortion (Hagler et al., 2019). T1-weighted images were acquired using a magnetization-prepared rapid acquisition gradient echo (MPRAGE) sequence (TR=2500, TE=2.88, flip angle=8) and T2-weighted images were obtained with fast spin echo sequence (TR=3200, TE=565, variable flip angle), with 176 slices with 1 mm^3^ isotropic resolution (Casey et al., 2018). Diffusion MRI data was acquired in the axial plane at 1.7 mm^3^ isotropic resolution with multiband acceleration factor 3. Ninety-six non-collinear gradient directions were collected with seven b0 images. Trained technicians inspected T1w, T2w, and dMRI images using a centralized quality control process in order to identify severe artifacts or irregularities (Hagler et al., 2019).

To assess gray matter macrostructure, we obtained baseline T1w and T2w images from the ABCD 3.0 release (NDA 3.0 data release 2020; https://dx.doi.org/10.15154/1520591) and implemented the Human Connectome Project minimal preprocessing pipeline (Glasser et al., 2013) at the Stevens Institute of Neuroimaging and Informatics. Regional parcellation and segmentation were then performed based on the Desikan-Killiany atlas in FreeSurfer 7.1.1 for each participant using T1w and T2w images (Desikan et al., 2006). The primary outcomes of interest included gray matter volume, thickness, and white matter volume in 68 cortical regions, the volume of 20 subcortical regions, as well as FA and MD for 19 white matter tracts (Hagler Jr. et al., 2009). For a complete list of regions by feature, please see Supplemental Table 1.

Tabulated white matter microstructure and demographic data from the baseline study visit were obtained from the 4.0 data release via the NIMH Data Archive (https://nda.nih.gov/abcd/; http://dx.doi.org/10.15154/1523041). ABCD diffusion processing employs five iterations of eddy current correction and robust tensor fitting to minimize gradient distortions and motion (Hagler et al., 2019; Hagler Jr et al., 2009). The b=0 images are coarsely registered to a diffusion atlas before being registered to T1w images via mutual information. DMRI images are then resampled and registered using the transform from rigid registration of the T1w image to the diffusion atlas. Finally, the diffusion gradient matrix is adjusted for head rotation. Probabilistic atlas-based tractography is then performed with AtlasTrack using a priori tract location probabilities to inform fiber selection (Hagler Jr et al., 2009). For this study, we utilized the tabulated FA and MD data from the AtlasTrack fiber atlas. Specifically, we selected the fornix, cingulate cingulum, parahippocampal cingulum, uncinate fasciculus, superior longitudinal fasciculus, inferior longitudinal fasciculus, inferior fronto-occipital fasciculus, anterior thalamic radiations, corticospinal tracts, and corpus callosum as regions of interest (ROIs).

### Analyses

All statistical analyses were conducted in R (R Core Team, 2019) with the vegan (Oksanen et al., 2022), lme4 (Bates et al., 2015), effectsize (Ben-Shachar et al., 2020), and bayestestR (Markowski et al., 2019) packages. We characterized variance and distributional overlap in brain outcomes between the sexes, investigated inhomogeneity of variance between the sexes with the Fligner-Killeen test, and implemented an analysis of similarities (ANOSIM). To ascertain whether variance, distributional overlap, and ANOSIM findings between the sexes were partially driven by additional variables, we repeated these analyses using residuals of brain outcomes after adjusting for additional variables (see details below).

For each ROI, we first compared the variance between group means to the within-sex variance for each ROI to determine whether the differences between sexes exceeded the differences within each sex for each ROI. To compare the within-sex variance of males and females for each ROI, we also calculated the coefficient of variation (CV), which accounts for potential scaling effects (Del Giudice, 2022). We then examined inhomogeneity of variance between males and females via the Fligner-Killeen test, which compares the variances of two groups using a median-centered chi-square χ ) test (Fligner & Killeen, 1976). We also calculated the overlap coefficient (OVL) for each ROI using the bayestestR package in R (Markowski et al., 2019), which measures the percentage of the sample that falls within the overlap between two distributions. To complement these descriptive analyses, we conducted an analysis of similarities (ANOSIM) with Euclidean distances with the vegan package in R (Oksanen et al., 2022). ANOSIM is a non-parametric method for comparing groups of a single sample on the basis of pairwise, ranked distances to determine whether the between-group differences are greater than the within-group differences. Significance is determined with a series of permutations that incrementally reorder group membership and calculate the proportion of permutations with an R greater than or equal to the observed R. ANOSIM R statistics range from -1 (all within-sex > between-sex ranked distances) to 1 (all between-sex > within-sex ranked distances) (also see Supplemental Table 2).

Residuals for each brain outcome were obtained from linear mixed modeling using the lme4 package in R (Bates et al., 2015; R Core Team, 2019). To account for additional sources of neuroanatomical variance beyond sex alone as well as site effects, the models included several independent variables as fixed effects along with data collection site as a random effect (i.e. the nesting of subjects within sites) (Supplemental Table 3). Age was measured in months and rounded to the nearest whole month. Pubertal development was assessed using the parent-report version of the Pubertal Development Scale (PDS) and categorized as prepuberty, early puberty, mid puberty, late puberty, and post-puberty (Cheng et al., 2021; Herting et al., 2020; Petersen et al., 1988; Thijssen et al., n.d.). Since few children in this age range were in late puberty or post-pubertal, we combined the mid, late, and post-puberty groups into a single category (mid/late puberty). We chose to include measures of race, ethnicity, and socioeconomic status in our models because human neurodevelopment is sensitive to various ecological factors which, due to systemic social injustice, are correlated with sociocultural variables, such as race, ethnicity, and socioeconomic status (Nketia et al., 2021; Werchan & Amso, 2017). Youth race was collected via caregiver report and caregivers were encouraged to select all answers that applied. Where more than one race was selected, we categorized participants as multiracial Black (if one of their selections was “Black”) or multiracial non-Black. Due to low group numbers, we combined Asian Indian, Chinese, Filipino/a, Japanese, Korean, Vietnamese, Other Asian, American Indian/Native American, Alaska Native, Native Hawaiian, Guamanian, Samoan, other Pacific Islander, and “other race” into a single category (“other race”). Youth ethnicity was parent-reported as either Hispanic or non-Hispanic. To encapsulate socio-economic status, we included educational attainment, operationalized as the highest level of education achieved in the household, and binned into the following categories: less than high school diploma, high school diploma or GED, some college, bachelor’s degree, or postgraduate degree. Idiosyncrasies of different MRI software and hardware can also impact brain segmentation (Liu et al., 2020), so we also included scanner manufacturer (Philips, Siemens, or GE) as a covariate. Lastly, we chose to include TBV as a covariate in our models of regional volume to account for the relationship between regional and whole-brain volume (Sanchis-Segura et al., 2020). Although studies of white matter microstructure generally do not adjust for whole-brain volume (Lebel et al., 2019; Takao et al., 2011), our recent findings in the ABCD cohort suggest adjusting for TBV can influence reported sex differences in FA and MD as well (Torgerson et al., 2024). Therefore, we elected to include TBV as a fixed effect but to conduct an additional set of white matter sensitivity analyses using the residuals without TBV in the models. TBV was calculated by FreeSurfer, then scaled by the sample root-mean-square.

## Results

A full description of the final sample for the current study can be found in Table 1. After stringent data cleaning, our final sample closely matched the full ABCD Study sample in terms of sex, age, pubertal development, ethnicity, and parental education but differed significantly in terms of race.

**Table 1.**
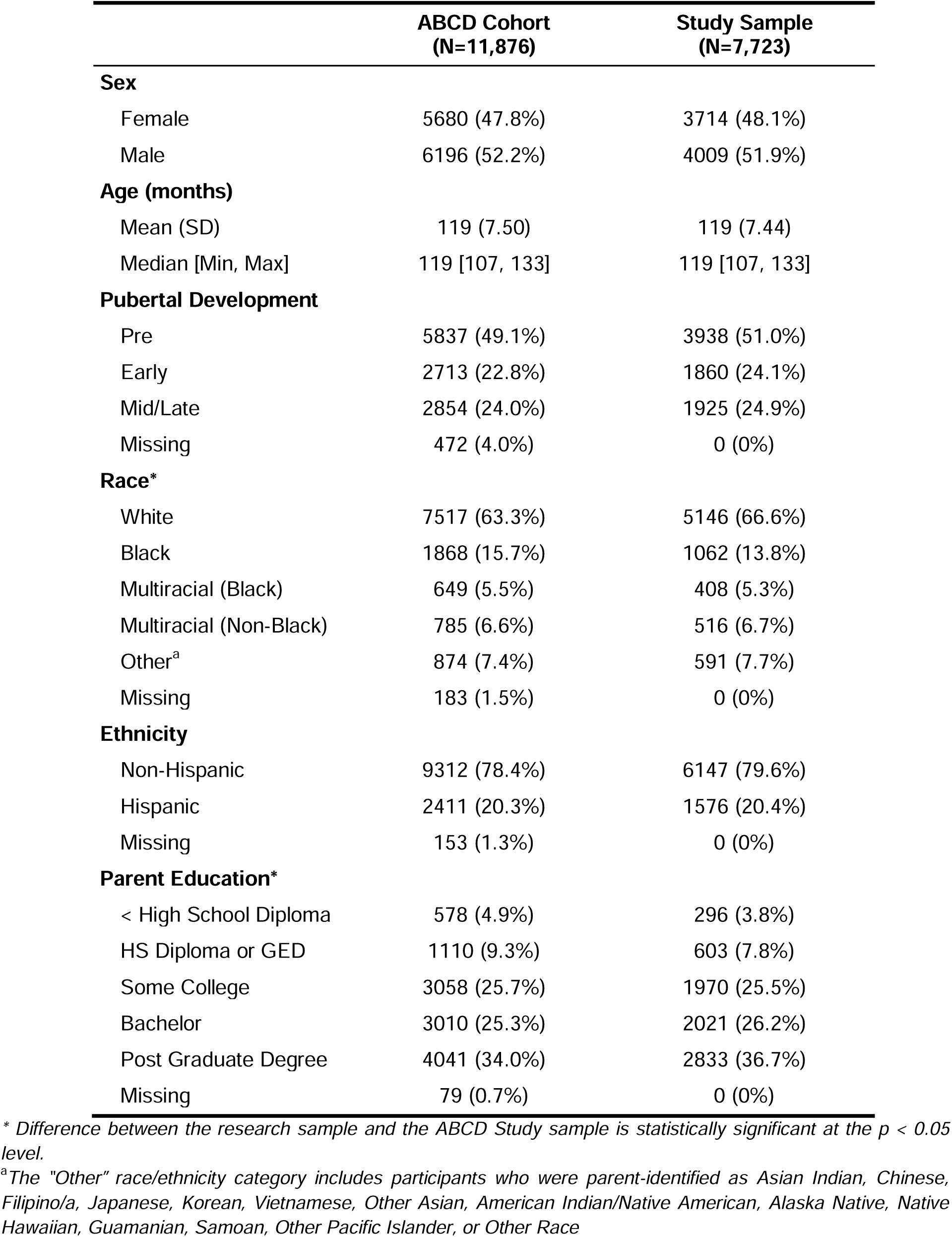
Demographic comparison between all ABCD Study subjects and the study sample.

### Global Brain Measures

In all global measures examined, within-sex variance exceeded between-sex variance, observed between the group means for male and female adolescents (Supplemental Table 4). For all whole-brain measures -both adjusted and unadjusted - the overlap between the male and female distributions was larger than the portions of the distribution unique to either sex (Figure 1). In the unadjusted data, we observed inhomogeneity of variance in TBV and white matter volume, such that male variance was greater than female variance, although the CV of unadjusted global measures were very similar between males and females (Supplemental Table 4). After adjusting for additional variables, inhomogeneity of variance was significant for TBV, white matter volume, mean FA, and mean MD (Supplemental Table 4). ANOSIM tests showed that within-sex and between-sex distances were similar for all whole-brain measures examined (as denoted by ANOSIM R < 0.1), with the exception of TBV and total white matter volume, which were similar with some differences (i.e., R < 0.25) (Figure 2; Supplemental Table 2). When adjusted values were used, all global measures showed similar variance between-and within-sex (Figure 2).

**Figure 1.**
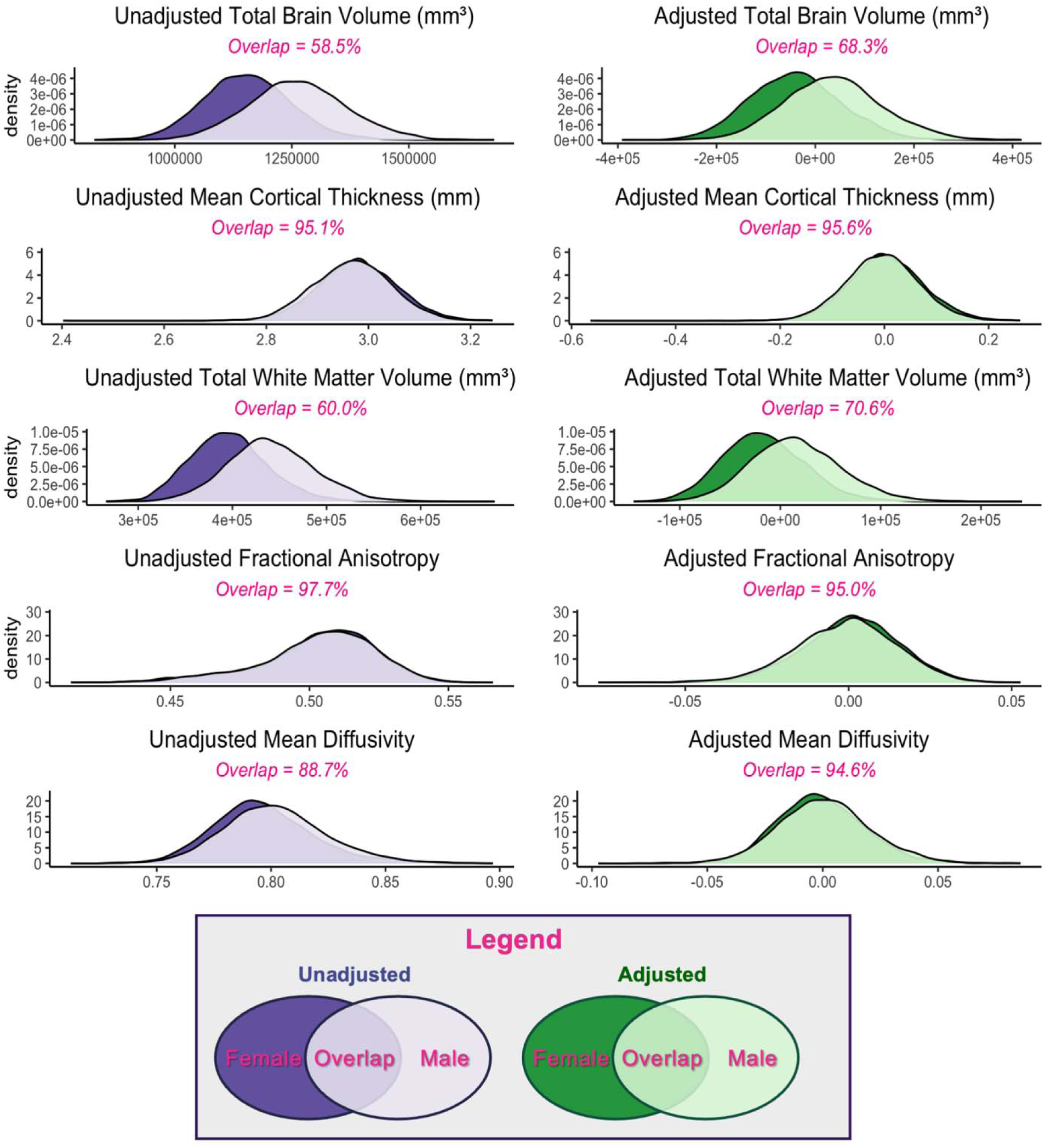
Overlap of global brain metrics in early adolescent males and females. Density plots and overlap of whole-brain measurements for both the unadjusted (purple) and adjusted (i.e., residual estimates, green) estimates for male (light) and female (dark) adolescents. Please note that the x-axis and y-axis change between measures (i.e. between brain volume and FA) due to large differences in scale.

**Figure 2.**
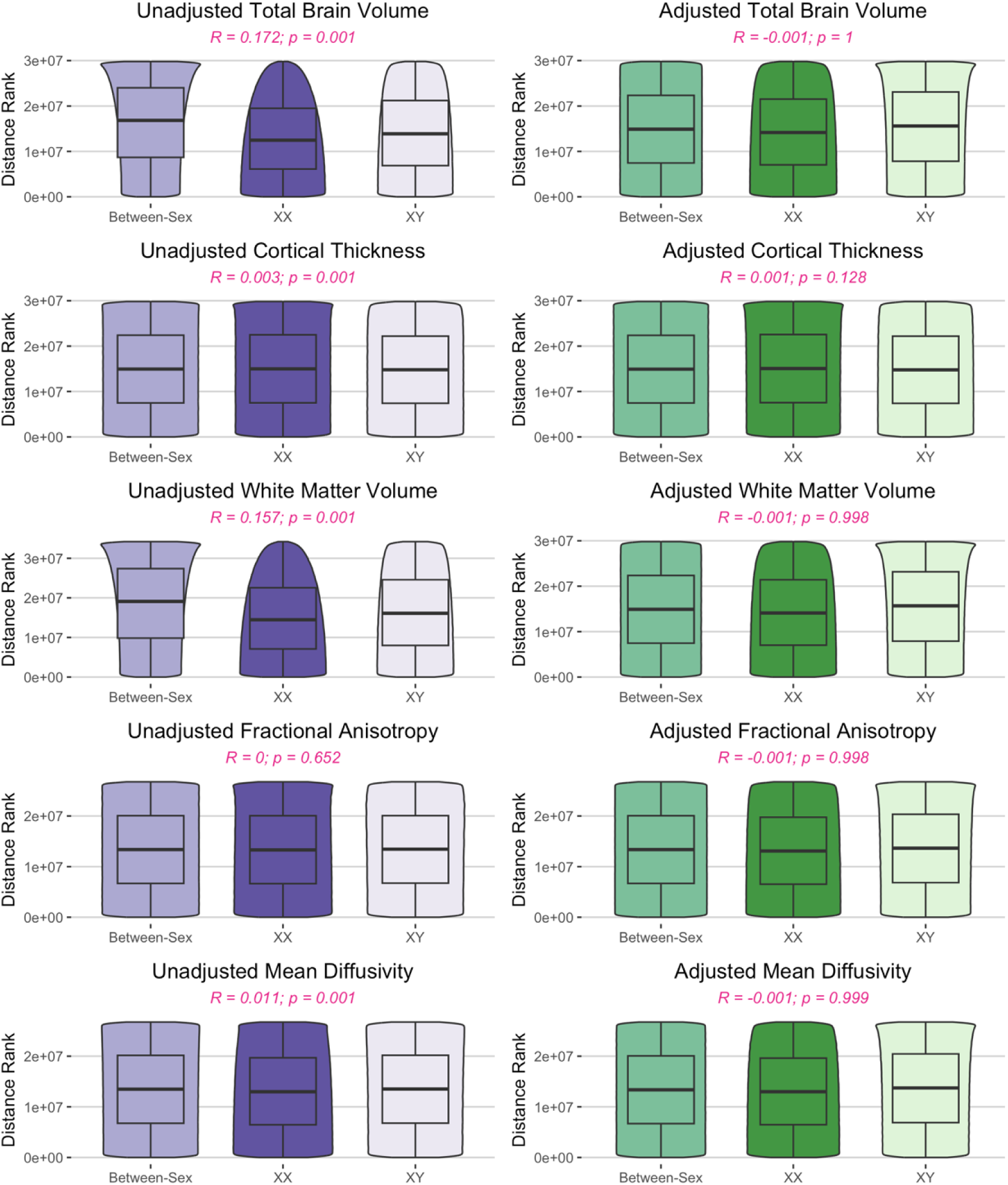
Similarity of within-and between-sex variance of global brain metrics in early adolescence. Violin plots of the within-sex and between-sex ranked distances from analysis of similarities (ANOSIM) test for both unadjusted (purple) and adjusted (residual estimates, green) as well as ANOSIM R statistic and FDR corrected p-values. An ANOSIM R statistic <0.1 suggests that group ranks are similar.

### Regional Gray Matter and Subcortical Macrostructure

The coefficients of variation for cortical volumes, subcortical volumes, and cortical thickness for male and female adolescents can be found in Supplemental Figures 2-4. In all cortical gray matter and subcortical volumes as well as cortical thickness regions examined, the variance between group means was smaller than the within-sex variance using both the unadjusted and adjusted volumes (Supplemental Tables 5-7). For cortical and subcortical volumes, inhomogeneity of variance between sexes was significant in all regions, with greater variance among male adolescents than among female adolescents (Figure 3A-B). The greatest sex differences in variance were seen in the supramarginal gyrus and central corpus callosum. In contrast to volume, female cortical thickness variance significantly exceeded male variance in the left superior frontal gyrus, left parahippocampal gyrus, and bilateral pericalcarine and lateral orbitofrontal cortices (Figure 3C). Similar to the whole-brain analysis, the overlap coefficients were also large for both the unadjusted and adjusted cortical gray matter volumes, subcortical volumes, and cortical thickness (Figure 4A-C). As expected, adjustment for TBV and other sources of variance led to an increase in the overlap of male and female regional cortical volumes (<u>unadjusted</u>: OVL range = 0.688 - 0.921, median = 0.788; <u>adjusted</u>: OVL range = 0.899 - 0.972, median = 0.939) and subcortical volumes (<u>unadjusted</u>: OVL range = 0.659 - 0.921, median = 0.749; <u>adjusted</u>: OVL range = 0.896 - 0.959, median = 0.939). Although the ANOSIM permutation tests were significant in 26 cortical and subcortical ROIs after FDR correction, the R statistic was consistently low (<u>unadjusted</u>: R statistic range: 0.0008 - 0.1171; median = 0.0446; <u>adjusted</u>: R statistic range: -0.0013 - 0.0086; median = 0.0001), indicating that the between-sex variance and within-sex variance were similar (Figure 5A-B). The same pattern was found in cortical thickness, where 56% of regions were significant before adjustment, albeit with R statistic values reflective of no meaningful difference in rank between the groups (<u>unadjusted</u>: R statistic range -0.0004 - 0.0194; median = 0.0013; <u>adjusted</u>: R statistic range -0.0007 - 0.0100; median = 0.0008) (Figure 5C).

**Figure 3.**
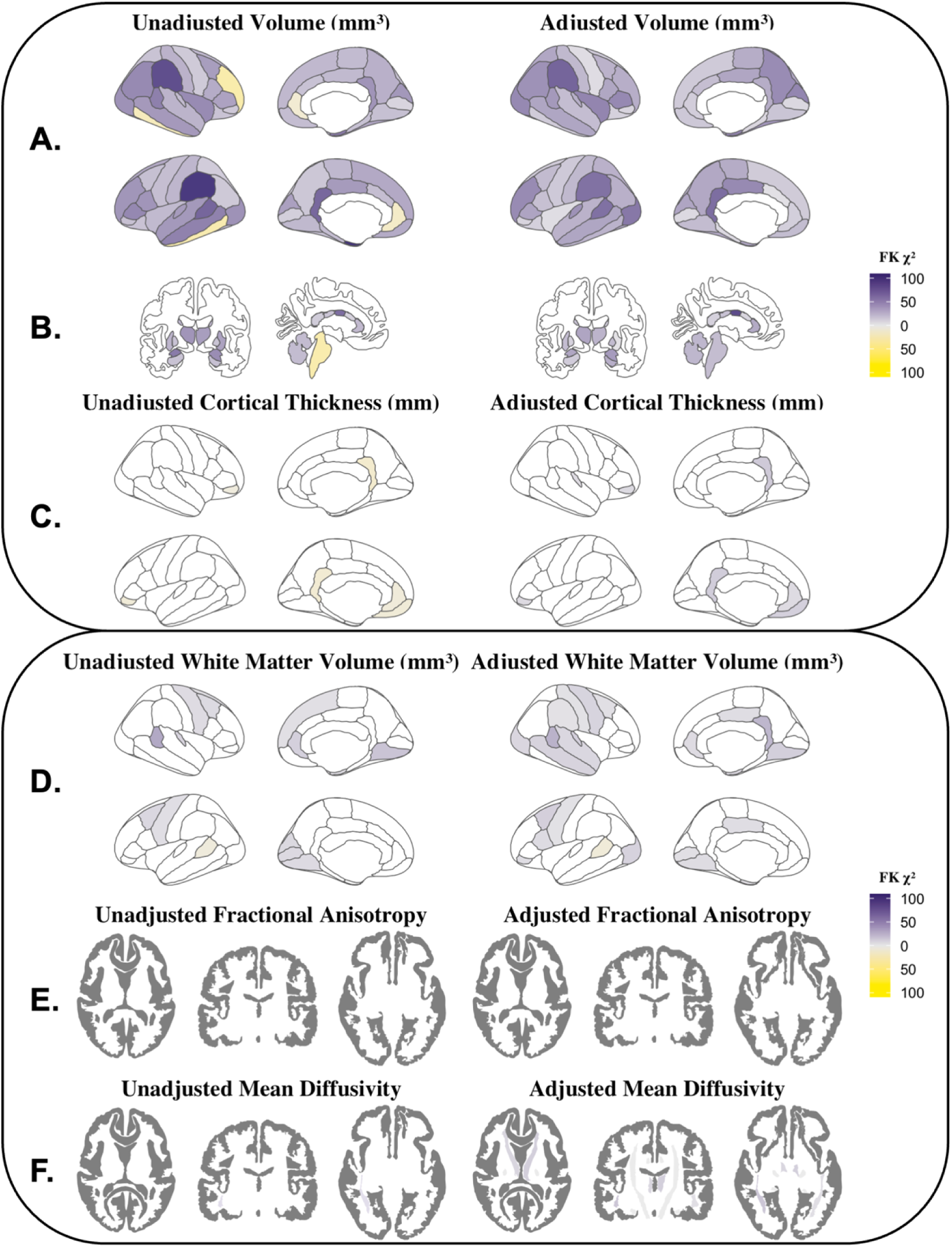
Unadjusted and adjusted inhomogeneity of variance between male and female adolescents for regional measures. A) cortical volumes, B) subcortical volumes, C) cortical thickness, D) white matter volumes, E) white matter fractional anisotropy (FA), and F) white matter mean diffusivity (MD). Colors reflect Fligner-Killeen χ2 statistic: purple denotes males > females; yellow denotes females > males.

**Figure 4.**
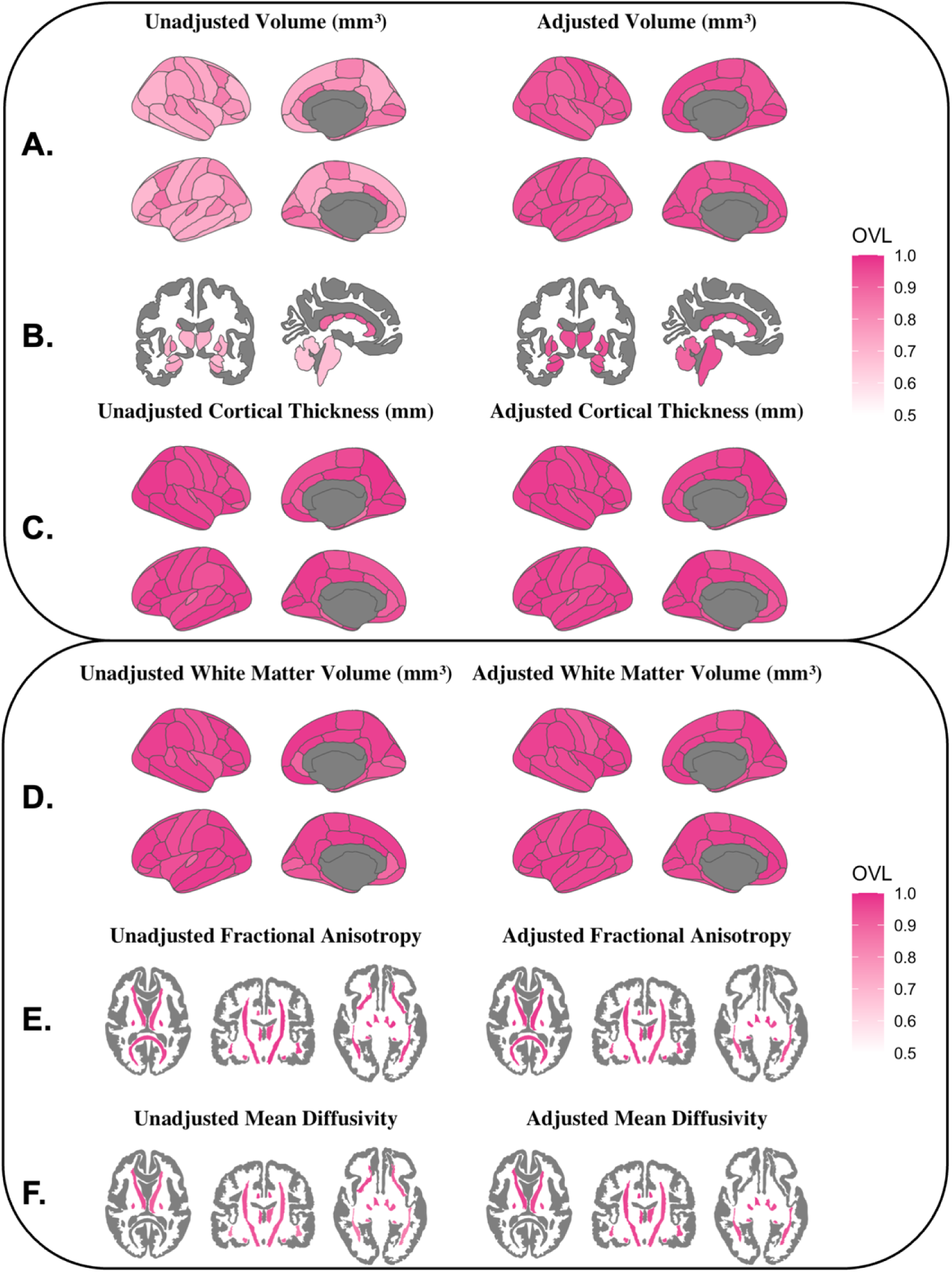
Unadjusted and adjusted overlap coefficients of male and female distributions for regional measures. A) cortical volumes, B) subcortical volumes, C) cortical thickness, D) white matter volumes, E) white matter fractional anisotropy (FA), and F) white matter mean diffusivity (MD). The overlap coefficient (OVL) compares the common area between two distributions to the unique variance and ranges from 0 (non-overlapping) to 1 (identical distributions).

**Figure 5.**
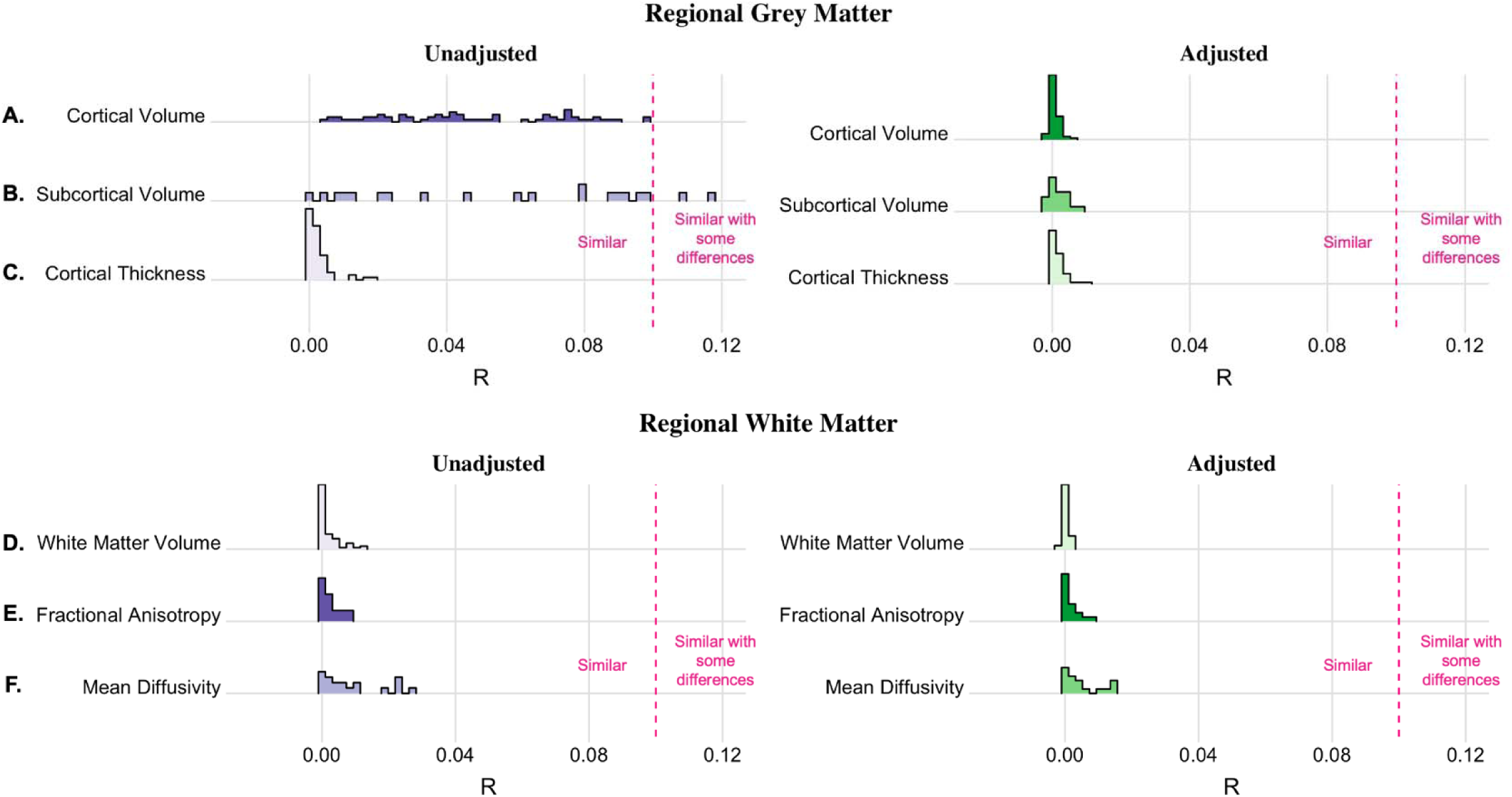
Distribution of Analysis of Similarities (ANOSIM) R statistics for unadjusted and adjusted regional measures. A) cortical volume, B) subcortical volume, C) cortical thickness, D) white matter volume, E) white matter fractional anisotropy, and F) white matter mean diffusivity. The ANOSIM R statistic ranges from -1 to 1, with 0 indicating no disparity in the magnitude of between-group and within-group pairwise comparisons. Please note that these figures are trimmed to increase visibility and therefore, the x-axis does not display the full range of possible R statistics. The dashed pink line denotes the threshold for groups to be considered “similar with some differences” (see Supplemental Table 2).

### Regional White Matter Volume

In all regions examined, the variance in white matter volumes between sexes was smaller than the within-sex variance (Supplemental Table 8, Supplemental Figure 5). Adjustment increased the percentage of regions with significant sex differences in variance (<u>unadjusted</u>: *p < 0.05* in 23.5% of regions; <u>adjusted</u>: *p < 0.05* in 41.2% of regions) (Figure 3D). Where significant, males showed greater regional variance than females except in the banks of the left superior temporal sulcus, where female variance exceeded male variance. Overlap coefficients were similar before and after adjustment (<u>unadjusted</u>: OVL range = 0.879 – 0.987, median = 0.963; <u>adjusted</u>: OVL range = 0.928 – 0.984, median = 0.963) (Figure 4D). The ANOSIM results were significant (*p* < 0.05) in 25/68 (37%) regions after FDR correction, yet the magnitude of the R statistic indicated that within-and between-sex variances were similar (<u>unadjusted</u>: R statistic range -0.0010 - 0.0132; median = 0.0002; <u>adjusted</u>: R statistic range -0.0011 - 0.0031; median = -0.0002; Figure 4D).

### White Matter Tract Microstructure

For both FA and MD the variance between the male and female mean values was universally smaller than the within-sex variance (Supplemental Tables 9-10; Supplemental Figures 6-7). In unadjusted FA values, no tracts showed significant inhomogeneity of variance. After adjusting for covariates, FA variance in the corpus callosum and right superior longitudinal fasciculus was significantly greater among male youth compared to female youth (Figure 3E). Before and after adjustment, male youth displayed significantly greater MD variance than female youth in 12/19 ROIs: the right corticospinal tracts, right uncinate fasciculus, corpus callosum, and bilaterally in the fornix, anterior thalamic radiations, superior longitudinal fasciculus, inferior longitudinal fasciculus, inferior fronto-occipital fasciculus, and superior longitudinal fasciculus (Figure 3F). Substantial overlap was observed (Figure 4E-F) in both the raw and adjusted FA (<u>unadjusted</u>: OVL range = 0.893 – 0.981, median = 0.959; <u>adjusted</u>: OVL range = 0.899 – 0.984, median = 0.963) and MD (<u>unadjusted</u>: OVL range = 0.829 – 0.967, median = 0.924; <u>adjusted</u>: OVL range = 0.928 – 0.977, median = 0.961). Similar to gray matter findings, the ANOSIM permutation tests found the ratio of between-sex to within-sex variance to be significantly different from 0 in many regions; however, the ANOSIM R statistic was small both before (<u>unadjusted</u>: R statistic range -0.0008 - 0.0267; median = 0.0027) and after adjustment (<u>adjusted</u>: R statistic range -0.0009 - 0.0156; median = 0.0018), suggesting similarity in rank distance between the groups (Figure 5E-F).

## Discussion

This study contextualizes previous reports of widespread group mean sex differences previously reported in early adolescence (Jamieson et al., 2023; Kurth et al., 2020; Lawrence et al., 2023; Lenroot et al., 2007; Peper et al., 2009; Raznahan et al., 2011b; Torgerson et al., 2024) by comparing the within-and between-sex variance as well as quantifying the neuroanatomical similarities between the sexes at ages 9 to 11 years old. In line with previous research in the developing brain (Bottenhorn et al., 2023; Forde et al., 2020; Wierenga et al., 2018), we detected significant inhomogeneity of variance between male and female youths. Moreover, we observed extensive overlap between male and female distributions and found between-sex and within-sex ranked differences to be similar in magnitude for all global and regional measures examined. We conclude that mean group sex differences in early adolescent brain structure are considerably smaller than the sex similarities, and therefore do not reflect distinct sex-based phenotypes (e.g., sexual dimorphism). Holistically, these results underscore the importance of accounting for within-group variance and inhomogeneity of variance when probing sex differences in brain morphology.

To assess similarity, we calculated the overlap (OVL) between male and female distributions in each global and regional measure. The OVL was invariably greater than 0.5, illustrating that across all structural metrics examined, more than half of all youths fell within the overlapping portion of the male and female distributions. In other words, there were substantial similarities between males and females throughout the brain. Similar results have been shown in adults, where “extensive overlap” has been reported between male and female distributions in all brain regions examined (Joel et al., 2015). While male and female total brain volume (TBV) distributions showed more similarity than difference (raw OVL = 0.585; corrected OVL = 0.682), TBV showed the least overlap between sex distributions of any measure examined, both before and after adjustment. This further supports its status as the largest and most replicable sex difference in pediatric brain structure (Ducharme et al., 2016; Lenroot et al., 2007; Levenstein et al., 2023; Paus et al., 2010; Sussman et al., 2016). However, brain size is related to overall body size (Burger et al., 2018; Schoenemann, 2004), so this difference may simply be a reflection of overall body size differences between male and female adolescents. Unadjusted regional overlap was lower for cortical and subcortical volume than for cortical thickness, FA, and MD -which had median regional OVLs greater than 0.9 before adjustment. After adjustment, overlap increased in most regions - particularly for regional volumes - and a minimum of 89.6% of the data fell within the overlap between male and female distributions for all adjusted regional measures. These findings further demonstrate that the brains of male and female youth appear very similar after accounting for additional sources of variance in the data. Therefore, our results extend the conclusions of Joel et al. (2015) to early adolescents and reaffirm that human brain macrostructure does not exist in binary, sexually dimorphic categories associated with sex, nor does it appear to exist on a continuum between male and female extremes.

This work expands upon previous findings of sex differences in within-sex variability in childhood (Bottenhorn et al., 2023; Wierenga et al., 2018). Wierenga et al. reported greater male variability in gray matter volume, whereas Bottenhorn et al. found greater male variability in white matter change over time, but greater female variability in cortical macro-and micro-structural change over time. After adjustment, we found significant sex differences in variance for TBV, average FA, average MD, and all regional volumes, with large inhomogeneity in the parietal lobe, basal ganglia, and limbic regions. Male variance exceeded female variance in all gray matter volume regions both before and after adjustment. Higher male variability in volume and diffusivity may be due, in part, to random X chromosome inactivation: heterozygous females express two different alleles of a single gene in a mosaic pattern throughout the brain, whereas homozygous females and males with a single X chromosome exhibit uniform expression (Raznahan et al., 2018; Raznahan & Disteche, 2021). Consequently, if two alleles of an X-chromosome gene have opposite effects, males and homozygous females will exhibit one of two extreme phenotypes, while heterozygous females will exhibit a mixed phenotype, decreasing the average trait variability among females. These results suggest that male structural variability is greater than female structural variability in gray matter volume and white matter microstructure, whereas female variability exceeds male variability in cortical thickness. Therefore, future research should examine the link between X-chromosome genes and regional gray matter volumes, while other sources of sex-related variance - such as estrogen and testosterone differences (Herting et al., 2015; Savic et al., 2017), BMI (Laurent et al., 2020), aerobic fitness (Chaddock-Heyman et al., 2015; Ruotsalainen et al., 2020), or eating behaviors (Breton et al., 2024) - should be explored with regard to cortical thickness variance.

Many univariate methods of comparison (i.e., t-tests, ANOVA) rely on the assumption of homogeneity of variance. Consequently, such tests are inappropriate for comparing sexes on measures with significant inhomogeneity of variance between sexes, such as gray matter volume. Given the combination of large within-sex variance and high overlap between distributions of male and female youth, it is important to instead test whether between-sex differences surpass within-sex differences. Thus, we used ANOSIM to assess the relative magnitude of all pairwise differences between subjects and test for significant differences between the within-group and between-group pairings. Although permutation tests indicated that in some regions we could reject the null hypothesis (i.e., within-sex and between-sex variances do not differ), it is possible for a statistical result to be “significantly different from zero yet inconsequentially small” in a sufficiently large sample (Dick et al., 2021; Warwick, 2001). For example, in the adjusted data, ANOSIM indicated that between-sex pairings were significantly different from within-sex pairings in 33% of ROIs, yet the maximum observed ANOSIM R statistic in the corrected regional data was 0.0156 (adjusted R range: -0.0013 - 0.0156). ANOSIM R statistics less than 0.1 indicate that the size of the difference between two adolescents of the same sex is similar to the size of the difference between two adolescents of the opposite sex (C. E. Arnold et al., 2021; Clarke & Gorley, 2001; Davis Birch et al., 2023; Sanchis-Segura et al., 2022). The fact that the results were significantly different from 0, but also very similar to 0 suggests that the sample size is sufficiently large to produce results with statistical significance but little practical or clinical significance. The ubiquity of the high overlap and low R statistic demonstrates that high similarity exists even in the measures with the highest mean sex differences. For instance, the effect size of sex for TBV (f^2^ = 0.243) would be considered medium-sized by Cohen’s standards (Cohen, 1992) and “extremely above average” for the ABCD dataset (Dick et al., 2021; Owens et al., 2021). Nonetheless, the TBV overlap was still greater than the difference (corrected OVL = 0.683) and the within-sex and between-sex differences were similar in size (corrected ANOSIM R statistic = 0.10). This highlights the fact that it is possible to have a relatively large, statistically significant sex effect even when subjects of the same sex differ about as much as subjects of different sexes. It is therefore critical for future analyses of sex to account for the mean-variance relationship and consider non-parametric methods that do not assume homogeneity of variance between sexes.

Taken together, these results contradict claims of sexual dimorphism in pediatric brain structure and contextualize the discussion of sex differences. This distinction between sexual dimorphism and sex differences is meaningful not just in theory, but also in practice. The putative sexual dimorphism of the developing brain has been cited in arguments for single-sex education (Bigler & Signorella, 2011; Eliot, 2013; Halpern et al., 2011) and as evidence in court cases regarding the rights of juveniles (Kennedy, 2021; *Re Alex: Hormonal Treatment for Gender Identity Dysphoria*, 2004). Yet, the large overlap between male and female distributions, small ratio of between-sex to within-sex differences, and significant inhomogeneity of variance reported here indicate that average pediatric sex differences are likely due to disparities in variability rather than two distinct phenotypes with a large mean difference. This lends credence to arguments that conventional methods for preclinical and clinical research of sex differences are not well-designed for application to personalized medicine and are insufficient to address health disparities between males and females (DiMarco et al., 2022; Miller et al., 2015; Richardson et al., 2015). Future research designs should employ more robust statistical methods and focus on precise sex-linked variables, such as hormones, chromosomes, gene expression, body size and composition, or social determinants of health.

### Limitations

Due to the cross-sectional nature of this study and the narrow age range of the participants, our results are limited in scope. As such, they should not be assumed to generalize to brain structure in early childhood, later in adolescence, adulthood or to longitudinal trajectories of brain development. Instead, they offer an in-depth look at the neuroanatomy of children between 9 and 11 years old. Furthermore, although sex is multifaceted and encompasses multiple hormonal, genetic, and gross anatomical features, we chose to focus on the presence or absence of a Y chromosome for our operational definition of sex. Consequently, it is unclear to what extent factors like hormone levels, gene expression, or X-chromosome inactivation play a role in our results. Additionally, as a non-experimental study, we cannot provide evidence of a causal link between sex chromosomes and variance. Since few studies examine the influence of social and environmental factors on neuroanatomical sex differences, some authors instead use the term “sex/gender” (Eliot et al., 2021; van Anders, 2022). While our previous work with data from the ABCD Study showed felt-gender did not explain a significant amount of variance in gray or white matter structure (Torgerson et al., 2024), we cannot rule out the possible influence of other sociocultural factors that may be correlated with sex.

Although this study discusses significance in terms of p-values (corrected for multiple comparisons), statisticians increasingly warn against dichotomous interpretations of results (i.e., “significant” or “nonsignificant”) (Gagnier & Morgenstern, 2017; Hoekstra et al., 2006) and overreliance on statistical significance to infer practical significance (Bangdiwala, 2016; Mohajeri et al., 2020). The frequency of small but significant f^2^ and ANOSIM R statistics found in this study further suggest that in such a large, diverse sample, p-values may not be reliable indicators of practical significance. This underscores the danger of dichotomous interpretation of statistical tests in large samples. As such, the significance of the inhomogeneity of variance results should also be interpreted with caution.

Moreover, the results may not be directly comparable between brain regions or metrics with very different mean outcomes (i.e. cerebellum volume vs. pars orbitalis volume, average cortical thickness vs. average FA). While this issue is frequently circumvented with standardization, we did not use this technique because it would have altered the variance we sought to characterize. Scaling was similarly rejected because of the associated reduction in significant digits for some measures. For example, when large values (such as TBV in mm^3^) are reduced to a smaller value (such as TBV in m^3^), the loss of precision could lead to more ties when rank-ordering the pairwise distances, ultimately impacting the ANOSIM results. Therefore, because of the regional differences in scale and the intrinsic link between the mean and variance, caution is urged when comparing results between different brain region outcomes.

### Conclusions

Early adolescent male and female brains are more similar than they are different. Due to high within-sex variability, the distributions of males and females have more overlap than difference on all measures of global and regional gray and white matter structure examined. Although male and female adolescents exhibited significant inhomogeneity of neuroanatomical variance, ANOSIM showed that within-sex and between-sex differences were similar in size. Overall, these results illustrate that sex differences in early adolescent brain structure do not amount to qualitative differences (e.g., sexual dimorphism), and that quantitative differences between sexes are likely too small to be practically meaningful compared with individual variability.

## Acknowledgements

Data used in the preparation of this article were obtained from the Adolescent Brain Cognitive Development^SM^ (ABCD) Study (https://abcdstudy.org), held in the NIMH Data Archive (NDA). This is a multisite, longitudinal study designed to recruit more than 10,000 children ages 9-11 and follow them over 10 years into early adulthood. The ABCD Study® is supported by the National Institutes of Health and additional federal partners under award numbers U01DA041048, U01DA050989, U01DA051016, U01DA041022, U01DA051018, U01DA051037, U01DA050987, U01DA041174, U01DA041106, U01DA041117, U01DA041028, U01DA041134, U01DA050988, U01DA051039, U01DA041156, U01DA041025, U01DA041120, U01DA051038, U01DA041148, U01DA041093, U01DA041089, U24DA041123, U24DA041147. A full list of supporters is available at https://abcdstudy.org/federal-partners.html. A listing of participating sites and a complete listing of the study investigators can be found at https://abcdstudy.org/consortium_members/. ABCD consortium investigators designed and implemented the study and/or provided data but did not necessarily participate in the analysis or writing of this report. This manuscript reflects the views of the authors and may not reflect the opinions or views of the NIH or ABCD consortium investigators. This study was also funded by RF1MH123223. We would like to acknowledge the assistance of Chun Chieh Fan, who provided the allele frequency ratios for participants in the ABCD Study.

