## Supplemental Figures for "More similarity than difference: comparison of within- and between-sex variance in early adolescent brain structure"

### Supplemental Figure 1.

*A graphical representation of exclusionary criteria and the quality control process for the study.*

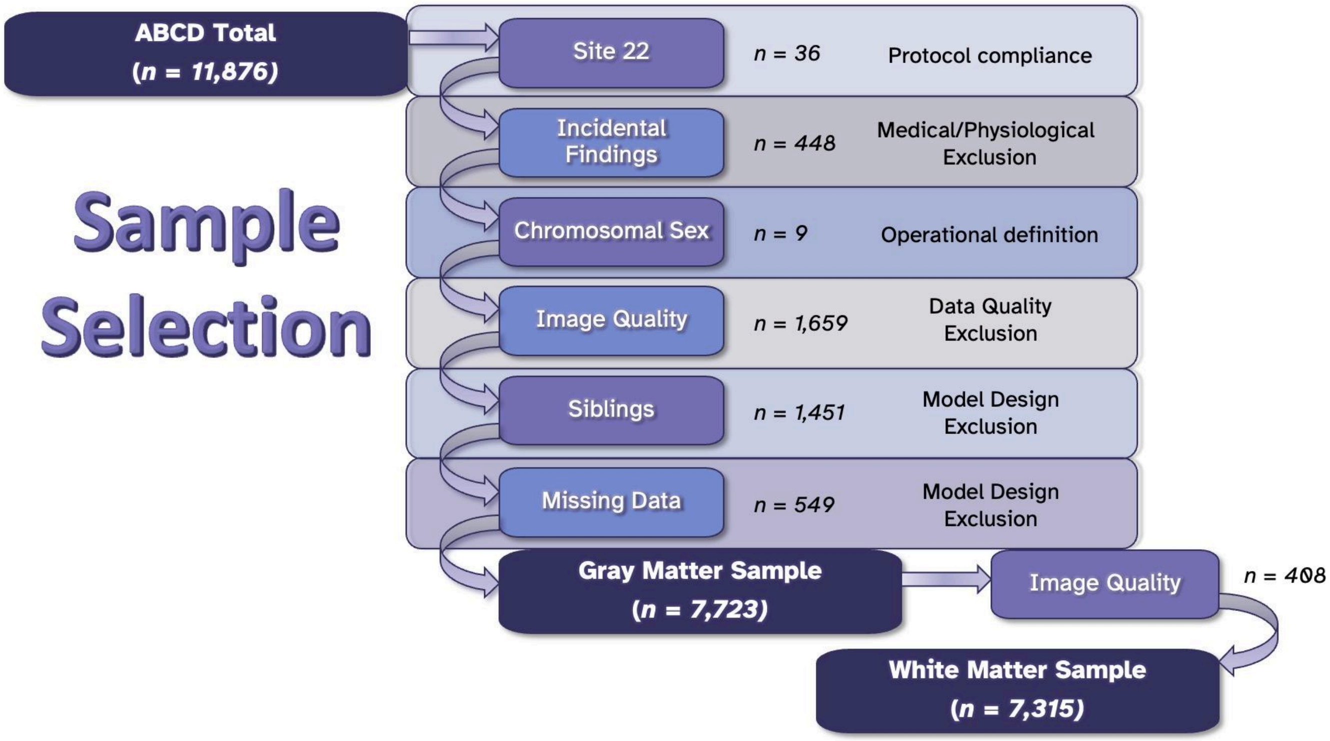

**Supplemental Figure 2.** ***Unadjusted coefficient of variance for males (blue) and females (pink) in regional cortical gray matter volume ROIs. * = inhomogeneity of variance FDR p-value < 0.05.***

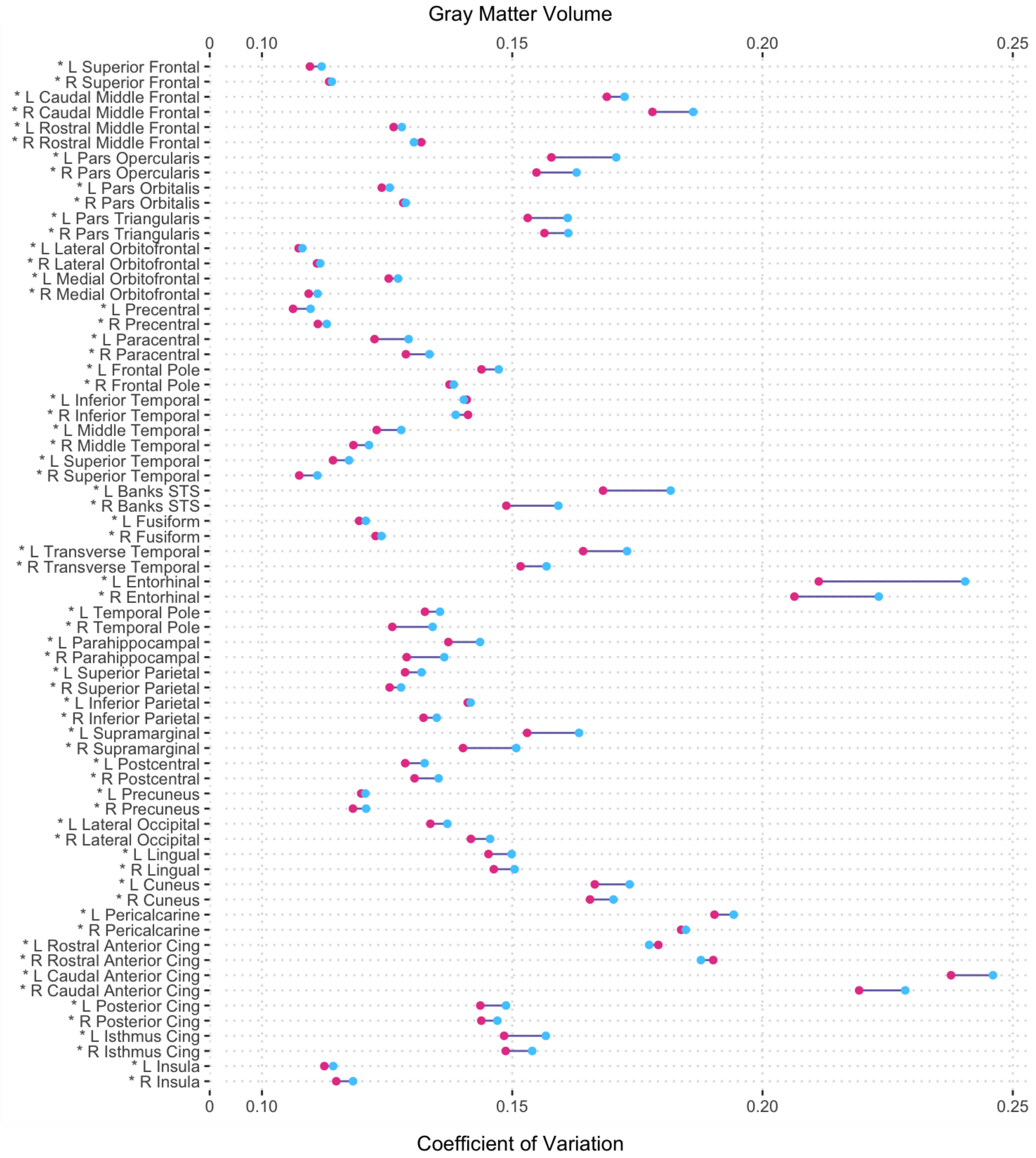

**Supplemental Figure 3.*****Unadjusted coefficient of variance for males (blue) and females (pink) in volumes for subcortical volume. * = inhomogeneity of variance p-value < 0.05.***

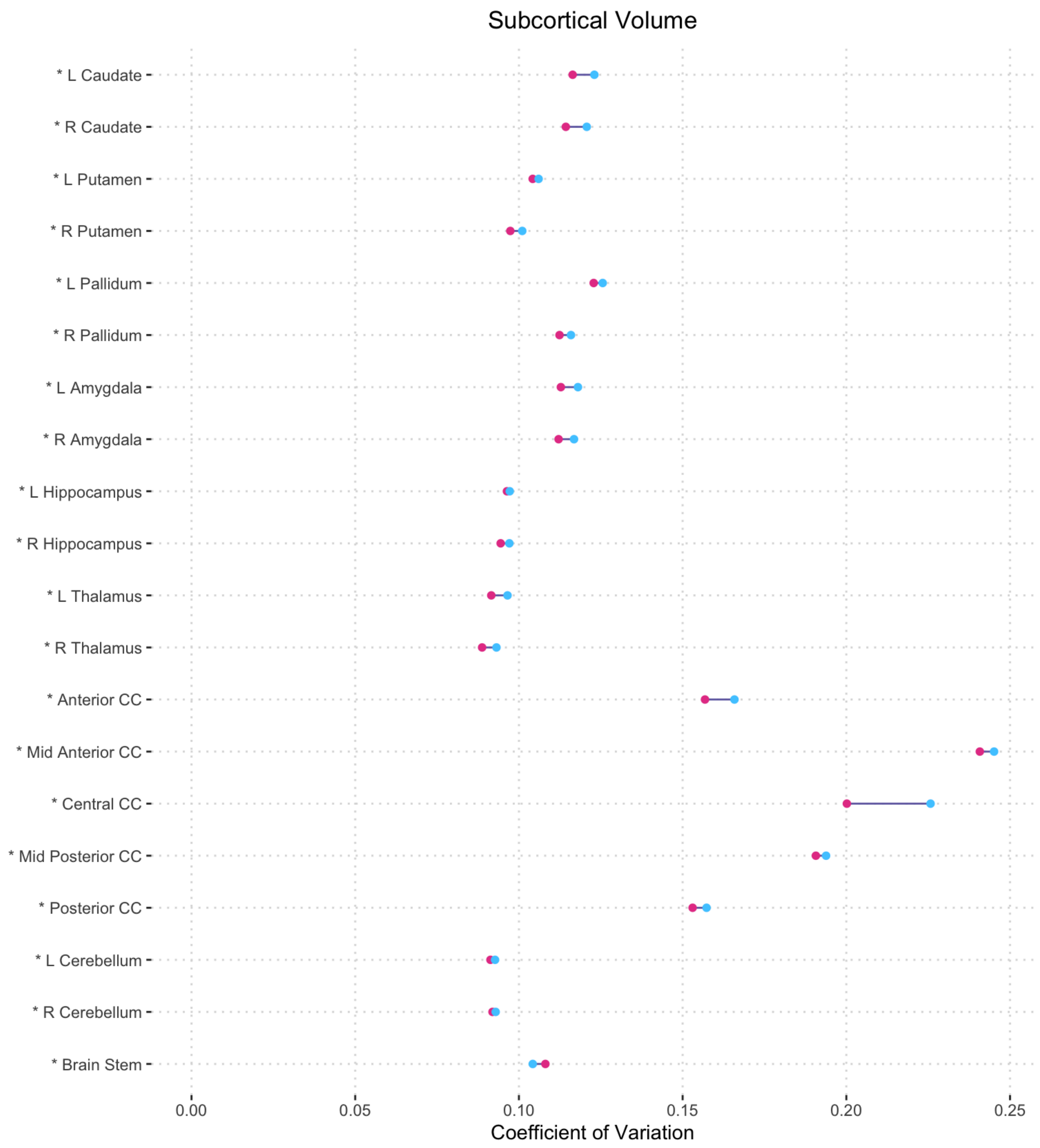

**Supplemental Figure 4.** ***Unadjusted coefficient of variance for males (blue) and females (pink) in regional cortical thickness. * = inhomogeneity of variance p-value < 0.05.***

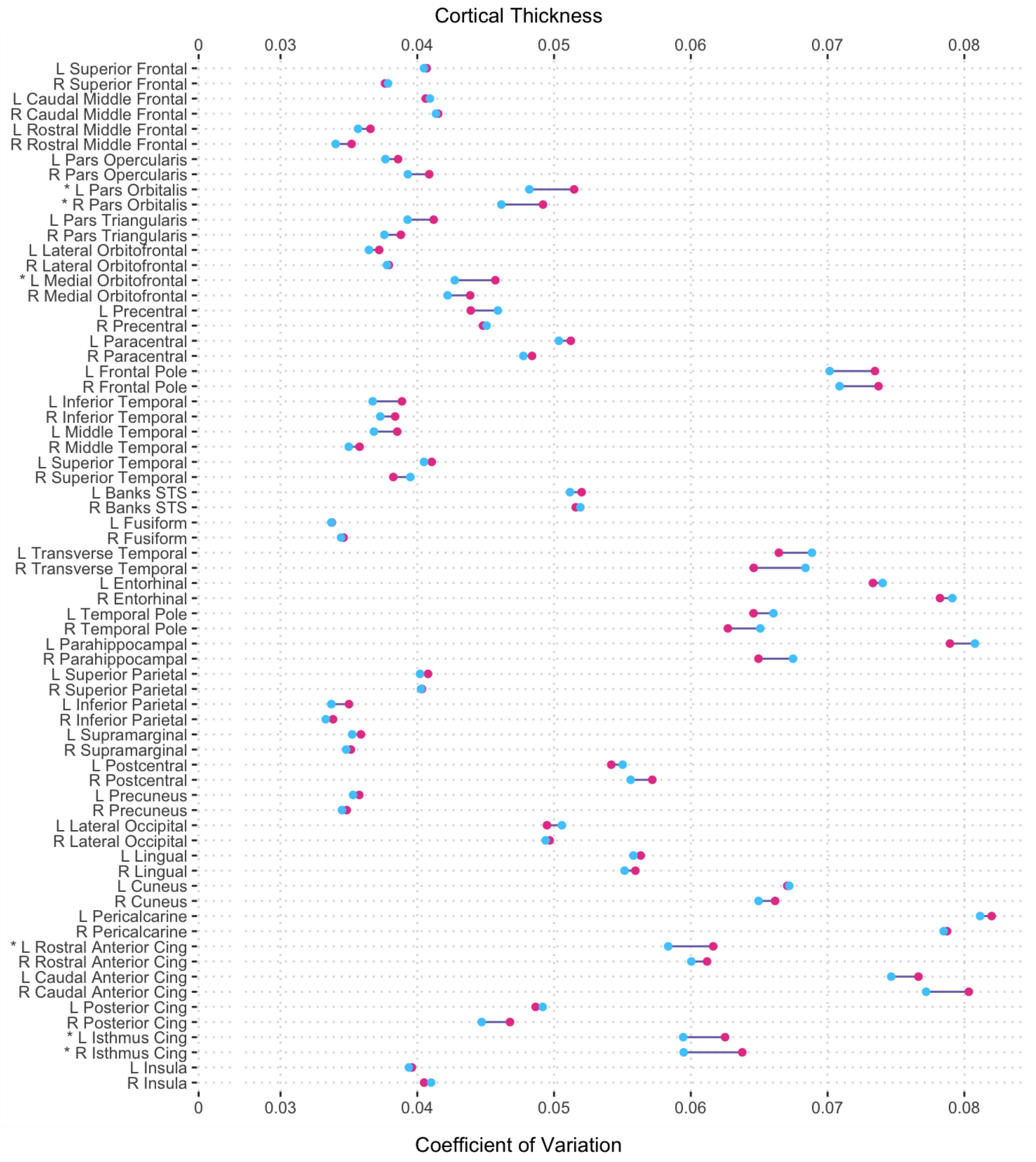

**Supplemental Figure 5.** ***Unadjusted coefficient of variance for males (blue) and females (pink) in regional white matter volume ROIs.*** *Abbreviations: L = left hemisphere, R = right hemisphere.* ***** denotes Fligner-Killeen $\chi^2$ test for inhomogeneity of variance FDR p-value < 0.05.

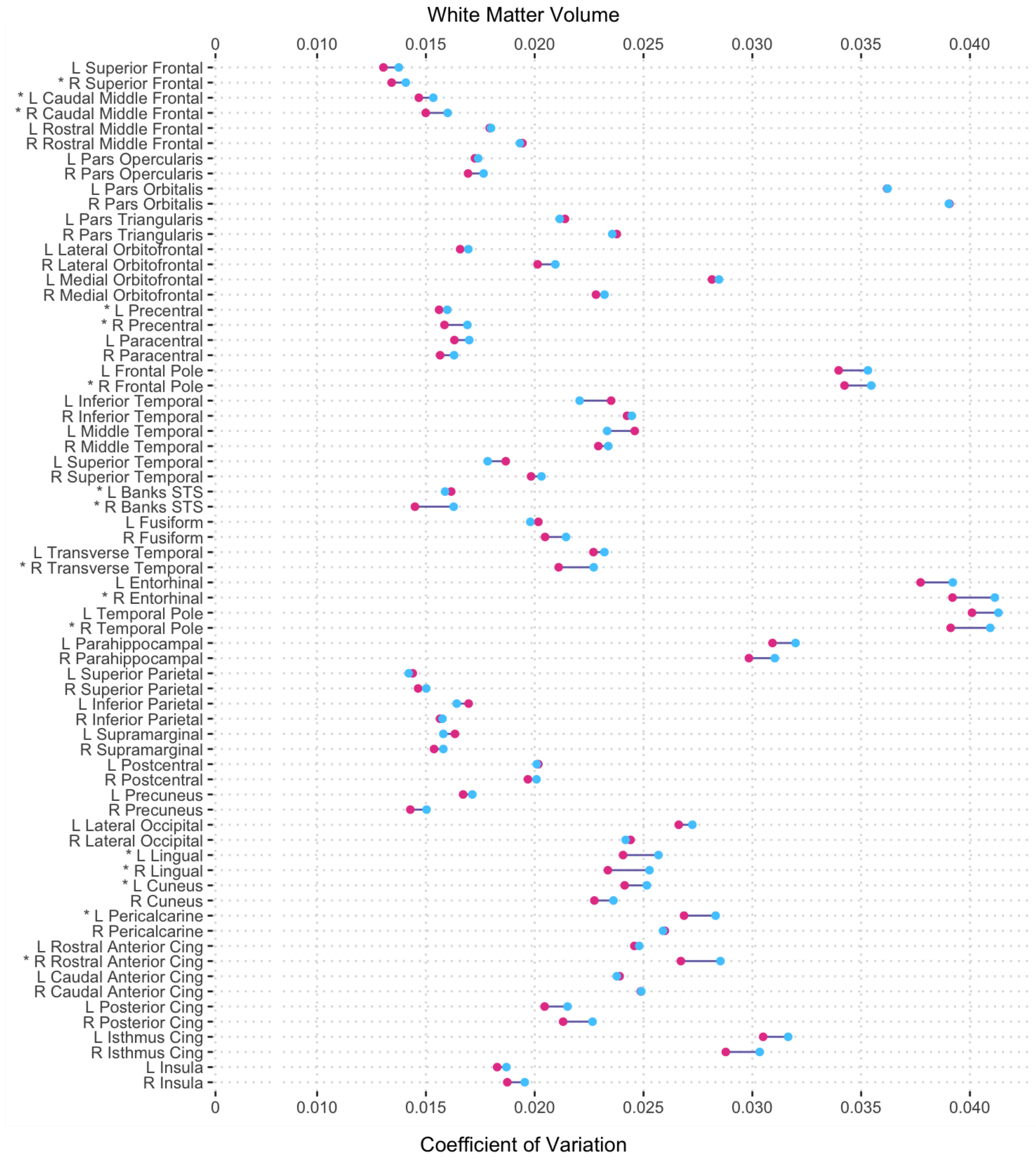

**Supplemental Figure 6.** ***Unadjusted coefficient of variance for males (blue) and females (pink) in regional FA.*** *Abbreviations: L = left hemisphere, R = right hemisphere, CST = corticospinal tract, ILF = inferior longitudinal fasciculus, IFOF = inferior fronto-occipital fasciculus.* ***** denotes Fligner-Killeen $\chi^2$ test for inhomogeneity of variance FDR p-value < 0.05.

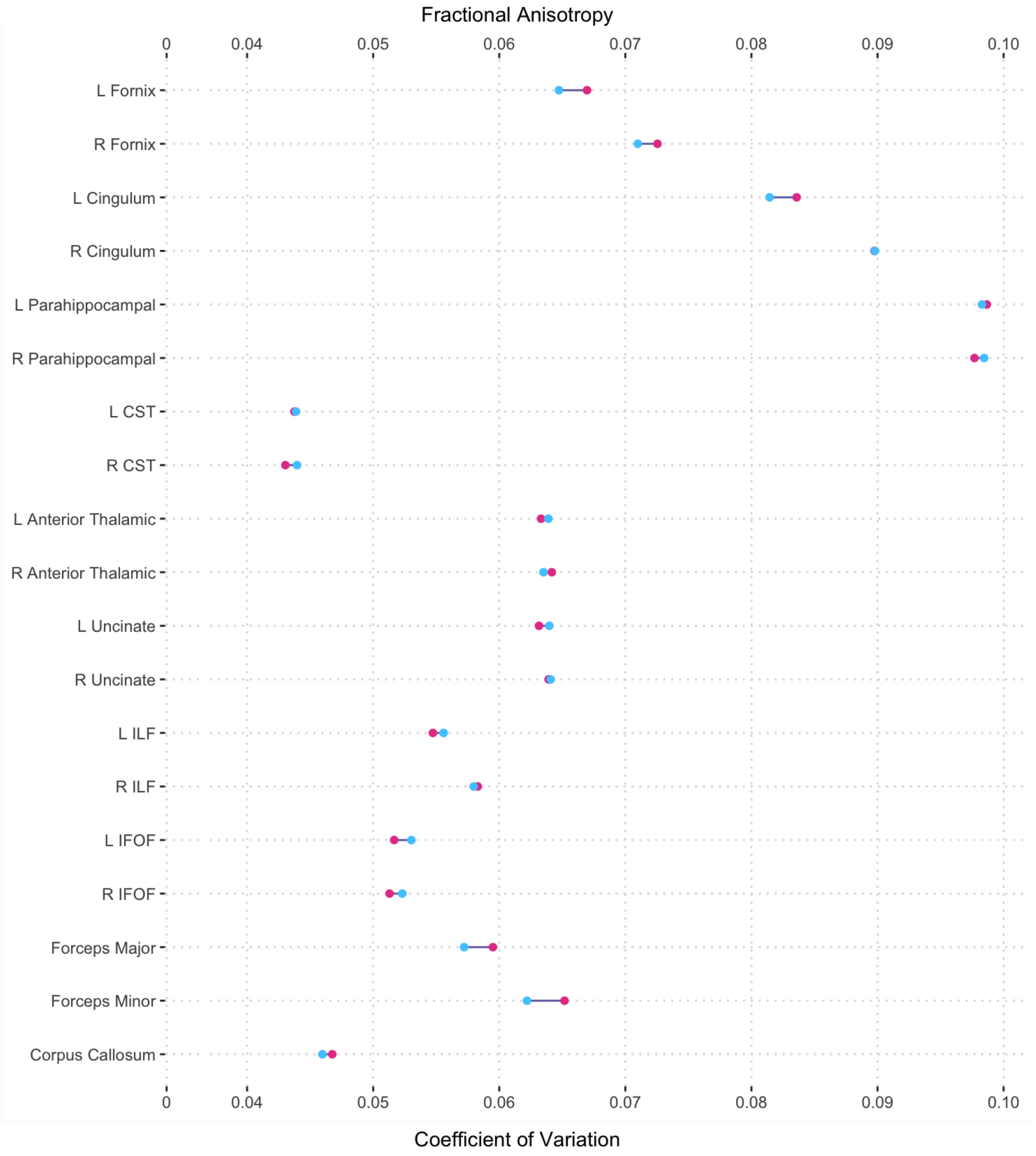

**Supplemental Figure 7.** ***Unadjusted coefficient of variance for males (blue) and females (pink) in regional MD.*** *Abbreviations: L = left hemisphere, R = right hemisphere.* ***** denotes Fligner-Killeen $\chi^2$ test for inhomogeneity of variance FDR p-value < 0.05.

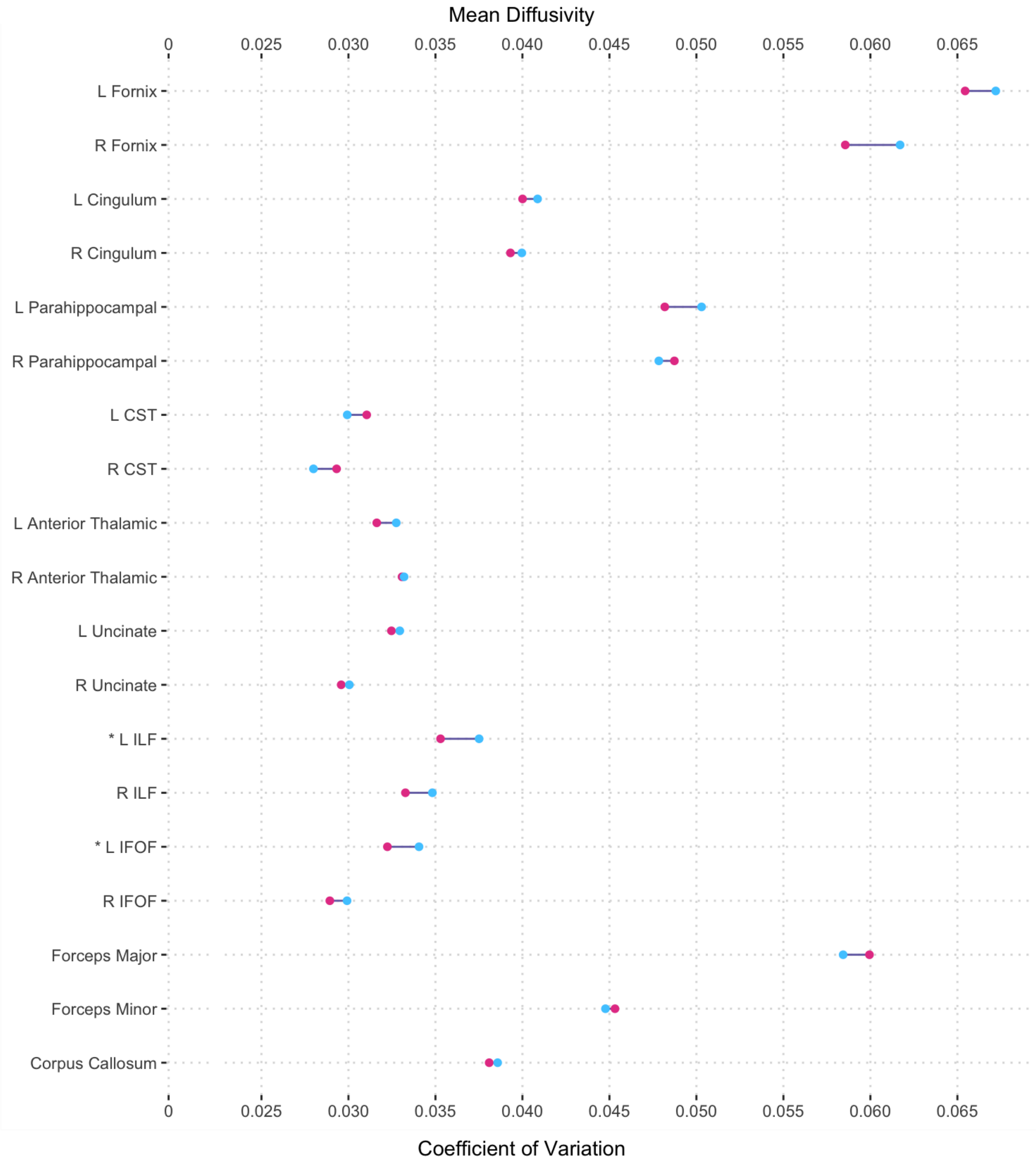
